# Direct evidence for decreased presynaptic inhibition evoked by PBSt group I muscle afferents after chronic SCI and recovery with step-training in the decerebrated rat

**DOI:** 10.1101/2020.05.05.078444

**Authors:** Guillaume Caron, Jadwiga N. Bilchak, Marie-Pascale Côté

**Author notes:** **Guillaume Caron, Ph.D.**, Drexel University College of Medicine, Marion Murray Spinal Cord Injury Center, Department of Neurobiology and Anatomy, Drexel University, Philadelphia, PA 19129. **Jadwiga Nichole Bilchak**, Drexel University College of Medicine, Marion Murray Spinal Cord Injury Center, Department of Neurobiology and Anatomy, Drexel University, Philadelphia, PA 19129. **Marie-Pascale Côté, Ph.D.** (corresponding author), Assistant Professor, Marion Murray Spinal Cord Injury Center, Department of Neurobiology and Anatomy, Drexel University, Philadelphia, PA 19129.

## Abstract

Spinal cord injury (SCI) results in the disruption of supraspinal control of spinal networks and an increase in the relative influence of afferent feedback to sublesional neural networks, both of which contribute to enhancing spinal reflex excitability. Hyperreflexia occurs in ~75% of individuals with chronic SCI and critically hinders functional recovery and quality of life. It is suggested to result from an increase in motoneuronal excitability and a decrease in presynaptic and postsynaptic inhibitory mechanisms. In contrast, locomotor training decreases hyperreflexia by restoring presynaptic inhibition.

Primary afferent depolarization (PAD) is a powerful presynaptic inhibitory mechanism that selectively gates primary afferent transmission to spinal neurons to adjust reflex excitability and ensure smooth movement. However, the effect of chronic SCI and step-training on the reorganization of presynaptic inhibition evoked by hindlimb afferents, and the contribution of PAD has never been demonstrated. The objective of this study is to directly measure changes in presynaptic inhibition through dorsal root potentials (DRPs) and its association to plantar H-reflex inhibition. We provide direct evidence that H-reflex hyperexcitability is associated with a decrease in transmission of PAD pathways activated by PBSt afferents after chronic SCI. More precisely, we illustrate that PBSt group I muscle afferents evoke a similar pattern of inhibition onto both L4-DRPs and plantar H-reflexes evoked by the tibial nerve in Control and step-trained animals, but not in chronic SCI rats. These changes are not observed after step-training, suggesting a role for activity-dependent plasticity to regulate PAD pathways activated by flexor muscle group I afferents.

**Key point summary:**

- Presynaptic inhibition is modulated by supraspinal centers and primary afferents in order to filter sensory information, adjust spinal reflex excitability, and ensure smooth movements.
- After SCI, the supraspinal control of primary afferent depolarization (PAD) interneurons is disengaged, suggesting an increased role for sensory afferents. While increased H-reflex excitability in spastic individuals indicates a possible decrease in presynaptic inhibition, it remains unclear whether a decrease in sensory-evoked PAD contributes to this effect.
- We investigated whether the PAD evoked by hindlimb afferents contributes to the change in presynaptic inhibition of the H-reflex in a decerebrated rat preparation. We found that chronic SCI decreases presynaptic inhibition of the plantar H-reflex through a reduction in PAD evoked by PBSt muscle group I afferents.
- We further found that step-training restored presynaptic inhibition of the plantar H-reflex evoked by PBSt, suggesting the presence of activity-dependent plasticity of PAD pathways activated by flexor muscle group I afferents.

## INTRODUCTION

Following chronic spinal cord injury (SCI), pathological adaptations lead to the development of spasticity and hyperreflexia in ~75% of individuals (Maynard *et al.*, 1990; Skold *et al.*, 1999; Holtz *et al.*, 2017). The symptoms range from hyperactive stretch reflex to increased muscle tone and from involuntary flexor withdrawal to extensor spasms (Nielsen *et al.*, 2007). These symptoms are the consequence of multiple mechanisms including the disengagement of supraspinal control below the injury site and subsequent increase in the relative influence of afferent feedback, hyperexcitability of motoneurons and decrease in spinal inhibitory mechanisms (Hultborn & Malmsten, 1983; Malmsten, 1983; Bennett *et al.*, 1999; 2001; Edgerton *et al.*, 2001; Boulenguez *et al.*, 2010; Brocard *et al.*, 2016). Increased activation of motoneurons by Ia afferents is thought to contribute to hyperactive stretch reflexes after SCI, which is a hallmark of spasticity. Historically, the hyperexcitability of the monosynaptic reflex (or H-reflex) after SCI has been suggested to rely on presynaptic inhibitory mechanisms (Thompson *et al.*, 1992; Calancie *et al.*, 1993; Faist *et al.*, 1994; Nielsen *et al.*, 1995; Schindler-Ivens & Shields, 2000; Morita *et al.*, 2001; Grey *et al.*, 2008). Presynaptic inhibition is a powerful mechanism that is under the control of supraspinal centers and sensory afferents. It selectively gates primary afferent transmission to spinal neurons by decreasing the probability of transmitter release from presynaptic terminals (reviewed in Rudomin & Schmidt, 1999; Rudomin, 2009). Classical presynaptic inhibition of the monosynaptic reflex was described by Eccles and colleagues more than half a century ago (Eccles, 1964) and shown to rely on GABAergic axo-axonic synapse onto sensory afferents. Presynaptic inhibition is paralleled by primary afferent depolarization (PAD) or its antidromic spread to dorsal roots as a dorsal root potential (DRP) which is mostly mediated by GABAergic receptors on sensory afferents and can be used as an estimate of its strength (Barron & Matthews, 1938; Eccles *et al.*, 1963a-b; Curtis *et al.*, 1971)

After acute SCI, tonic supraspinal inhibition of spinal reflex pathways is impaired, leading to disinhibition of spinal reflexes (Eccles & Lundberg, 1959; Carpenter *et al.*, 1963; Quevedo *et al.*, 1993). While the strong regulatory effect of supraspinal pathways on presynaptic inhibition suggests that it is significantly disrupted after chronic SCI, the reorganization of afferent control on presynaptic inhibition when supraspinal inhibition is disrupted remains to be determined. The first objective of this study is to directly investigate the changes in transmission in PAD pathways originating from hindlimb afferents after a chronic SCI through L4-DRP recordings and assess if a reduction in the compound PAD amplitude parallels the decrease in presynaptic inhibition of the H-reflex.

After chronic SCI and other conditions that lead to long-lasting immobilization, spinal reflex modulation is improved by locomotor training (Côté *et al.*, 2003; Côté & Gossard, 2004; Martin Ginis & Latimer, 2007; Knikou & Mummidisetty, 2014; Caron *et al.*, 2016). Activity-dependent recovery relies on sensory feedback during stepping as descending modulation is disrupted (Edgerton *et al.*, 2001). Increased phase-dependent modulation of sensory input with locomotor training after chronic SCI has been hypothesized to rely on the recovery of presynaptic inhibition for years (Bouyer & Rossignol, 2003; Côté *et al.*, 2003). In agreement with this putative return of presynaptic inhibition, the hyperexcitability of the monosynaptic reflex (or H-reflex) is decreased in both SCI humans and animals that followed a step-training regimen (Côté *et al.*, 2003; Knikou & Mummidisetty, 2014). The second objective of this study is therefore to directly investigate if step-training restores primary afferent control of presynaptic inhibition of the H-reflex in parallel to transmission in PAD pathways.

We hypothesized that 1) a decrease in PAD evoked by primary afferents originating from the hindlimbs contributes to the decrease presynaptic inhibition of the H-reflex after chronic SCI and 2) the beneficial effect of step-training on the recovery of presynaptic inhibition of the H-reflex is associated with a concomitant return of the PAD after chronic SCI. We used a decerebrated rat preparation to preserve spinal neuron excitability and to evaluate presynaptic inhibition evoked by PBSt on tibial afferents. We found that after chronic SCI, the compound PAD evoked by PBSt (posterior biceps – semitendinosus) and recorded as a DRP from a cut L4 dorsal rootlet is decreased. We simultaneously measured the L4-DRP and the plantar H-reflex evoked by the tibial nerve and tested the effect of a PBSt conditioning stimulation. We show that there is a decrease in transmission in PAD pathways activated by PBSt afferents, which parallels presynaptic inhibition of the plantar H-reflex, and contributes to hyperreflexia after chronic SCI. The recovery of presynaptic inhibition of the plantar H-reflex evoked by PBSt group I afferents with step-training further suggests a better modulation of tibial Ia afferents by PBSt. Because flexor muscle proprioceptors affect the activity of hindlimb muscles and exert a powerful control on locomotor rhythm (Grillner & Rossignol, 1978; Rossignol *et al.*, 2006), this suggests a contribution of activity-dependent plasticity in decreasing hyperreflexia and improving walking ability through restoration of presynaptic inhibition.

## METHODS

### Ethical approval

All procedures complied with ARRIVE guidelines, were performed in accordance with protocols approved by Drexel University Institutional Animal Care and Use Committee (IACUC) and followed the National Institutes of Health guidelines for the care and use of laboratory animals. Rats were housed in facilities accredited by AAALAC (Association for Assessment and Accreditation of Laboratory Animal Care) on a 12h light-dark cycle, with controlled room temperature, and ad libitum access to food and water. Rats were singly housed for 3 days following injury, then in pairs for the remainder of the study.

### Surgical procedures and postoperative care

Adult female Sprague Dawley rats (240-300g, Charles Rivers Laboratories) underwent a complete spinal transection at the low thoracic level (T12) as described previously (Côté *et al.*, 2011; Côté *et al.*, 2014). Briefly, rats were anesthetized with isoflurane (~2.5%) in O2 and a laminectomy was performed at the T10–T11 vertebral level under aseptic conditions. The dura was carefully slit open, the spinal cord completely transected with small scissors, and saline-soaked gel foam inserted in the cavity to achieve hemostasis. The completeness of the spinal cord lesion was confirmed by the retraction of the rostral and caudal portions of the cord and by examining the ventral floor of the spinal canal. Paravertebral muscles were sutured back together, and the skin closed with wound clips. Upon completion of the surgery, animals received a single injection of slow release buprenorphine (0.05 mg/kg, *s.c*.) and daily saline (5 ml, *s.c*.) and Baytril (100 mg/kg, *s.c*.) for 7 days to prevent dehydration and infection, respectively. Bladders were expressed manually at least twice daily until voiding reflex returned. Rats were randomly assigned to one of the following groups: chronic spinal cord injured (SCI; n=7) or chronic spinal cord injury + step-training (SCI + Step-training; n=6) with the terminal experiment taking place ~8 weeks postinjury. Another group of animals (Control; n=8; 260-460g) were acutely transected at the same spinal level at least 3h before starting terminal experiment recordings.

### Exercise regimen

Beginning on day 5-7 post-injury, the exercised groups received 10 min of daily exercise, 5 days per week until the terminal experiment. Animals undergoing step-training were placed on a treadmill belt (11-15cm/s) with the forelimbs resting on an acrylic glass platform. Perineal stimulation was provided to induce and maintain locomotor movements (Etlin *et al.*, 2010). Weight support was provided by the experimenter and adjusted at a level sufficient to prevent collapse of the locomotion until the animals recovered weight-bearing stepping (Côté *et al.*, 2011).

### Terminal experiments

Isoflurane anesthesia (~3.5%) in O2 was induced by mask and was maintained through tracheotomy during surgery. One common carotid artery was cannulated to monitor blood pressure while the other one was ligated. A cannula was inserted in a jugular vein for administration of Ringer-Locke solution (2.5% dextrose) to maintain the mean arterial blood pressure (>80 mmHg). The posterior biceps and semitendinosus nerves (PBSt) were dissected free from surrounding tissue and cut distally, while the tibial nerve was dissected free from surrounding tissue and left intact. A laminectomy was performed from T13 to L3 to expose the lumbar enlargement of the spinal cord and to allow appropriate identification of dorsal roots. Skin flaps surrounding the spinal cord and peripheral nerves were used to construct mineral oil pools. At this stage, a precollicular decerebration was performed, the rostral tissue was removed by suction and residual bleeding was prevented by packing the cranial fossae with small pieces of gelfoam soaked in cold thrombin solution. Anesthesia was then slowly discontinued, and rats artificially ventilated to maintain the expired CO2 near 4%. Animal rectal temperature was controlled and kept at 37°C via a DC heating pad and heating lamp. Recordings started at least 60 min after isoflurane was discontinued to ensure complete anesthesia removal and animal stabilization (Marchenko *et al.*, 2002).

### Stimulation and recordings

The experimental setup is illustrated in **Figure 1A**. PBSt and tibial nerves were mounted on bipolar hook electrodes (FHC, Bowdoin, ME, USA) for stimulation. The cord dorsum potential (CDP), elicited by the stimulation of peripheral nerves, is used to record the incoming volley with a monopolar silver ball electrode placed at the L4-L5 dorsal root entry. The stimulation threshold for the most excitable afferent was defined as the minimum intensity capable of producing an incoming volley as recorded from the CDP. The stimulation strength is thereafter expressed as multiples of this threshold (xCDP). Dorsal root potentials (DRPs) were recorded from cut L4 dorsal rootlet using a bipolar tungsten hook electrode, placed 1mm from the cord, in response to a stimulation (200 μs duration) of various intensities to the tibial or PBSt nerves. CDP and DRP recordings were amplified (x1000) and band-pass filtered (0.1 Hz-5 kHz; A-M System, Carlsborg, WA, USA). H-reflexes were evoked by a stimulation (200 μs duration) to the tibial nerve and recorded using bipolar wire electrodes (Cooner Wire, Chatsworth, CA, USA) inserted in the interosseous muscles. EMG recordings were amplified (x1000) and band-pass filtered (10 Hz-5 kHz; A-M Systems). All signals were digitized (10 kHz) and fed to Signal 5 software (Cambridge Electronic Design, UK).

**Figure 1.**
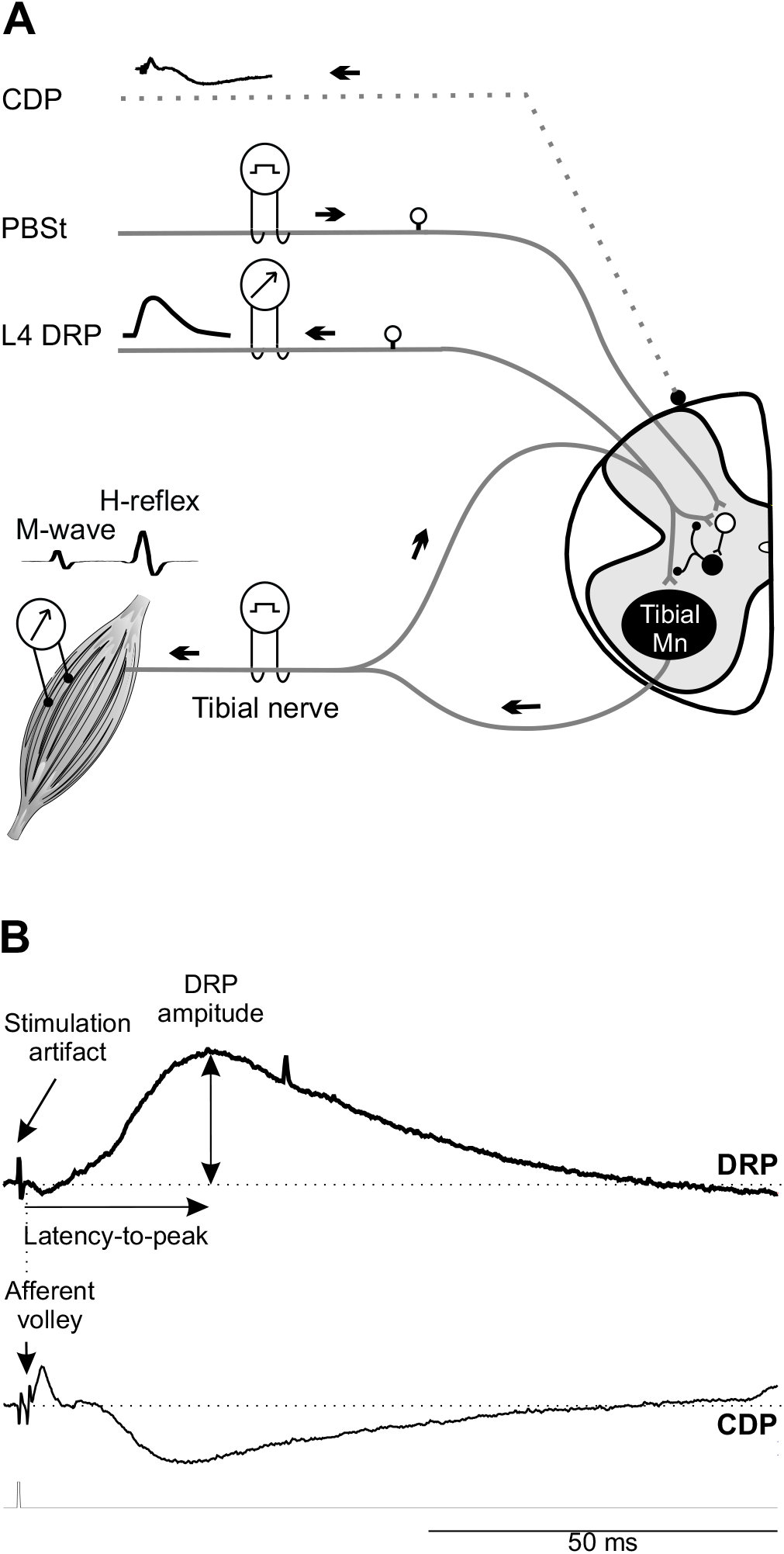
Experimental set-up. ***A*,** The PBSt nerve, tibial nerve and cut dorsal rootlet were mounted on bipolar hook electrodes. The cord dorsum potential (CDP) was recorded using a monopolar silver ball electrode placed at the L4-L5 dorsal root entry to determine the activation threshold for the most excitable afferent fiber. Dorsal root potentials (DRPs) were recorded in response to a stimulation to the tibial or PBSt nerves. H-reflexes were evoked by a stimulation to the tibial nerve and obtained using bipolar wire electrodes inserted under the plantar surface of the foot. In the conditioning protocol, PBSt stimulation was used as the conditioning stimulus and preceded the tibial stimulation. ***B***, Typical DRP recording displaying a long-lasting upward deflection. By convention, DRP negativity is represented upwards and CDP negativity downwards. The latency-to-peak was determined as the time between the onset of the afferent volley (CDP) and the maximal DRP amplitude from baseline to peak.

To determine the input-output relationship, PBSt- and Tib-evoked DRPs were recorded in response to stimuli of increasing intensity until reaching DRP_max_. M-waves and H-reflexes were also recorded in response to stimuli of increasing intensity to build a recruitment or input-output curve. Presynaptic inhibition evoked by group I afferents from PBSt on DRPs and H-reflex evoked by a stimulation to the tibial nerve were estimated using a conditioning-test protocol. Tibial nerve stimulations (1p, 1.2-1.8T) were alternatively preceded (conditioned) or not (test) by stimulation of PBSt (4p, 1.5T, 250 Hz) with conditioning-test (C-T) intervals ranging from 0 to 150ms. Each C-T combination was delivered at 0.2Hz.

### Data analysis

A typical DRP trace is shown in **Figure *1B***. The stimulation artifact is followed by a short downward deflection. The subsequent long-lasting upward response, representing the DRP, is clearly distinguishable. We measured the latency-to-peak as determined by the time between the onset of the afferent volley (cord dorsum potential, CDP) and the maximal amplitude of a given DRP. The DRP amplitude was measured and plotted as a function of stimulus intensity to build an input-output curve for each animal. The stimulation intensity required to reach the maximal DRP amplitude (DRP_max_) was then determined. A Boltzmann sigmoid function was then fitted to the data (Klimstra & Zehr, 2008; Lundbye-Jensen & Nielsen, 2008a; Smith *et al.*, 2015), and the goodness of the fit (R^2^) determined.

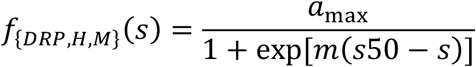

The estimated parameters of the function include the maximal amplitude (*a*_max_), the slope of the function (*m*) which reflects the gain independently from the absolute amplitude (Carroll *et al.*, 2001), and the stimulation intensity required to elicit a DRP equivalent to 50% of DRP_max_ (s50) at a given stimulation intensity (*s*). Group averages were calculated from the individual values for *a*_max_, *m* and s50, and used to establish a sigmoid function for each group. Similarly, we also performed measurements of the input-output curve for the M-wave and H-reflex for each animal. The motor threshold, H-reflex threshold, maximal H-reflex and M-wave amplitude and latency were also determined.

To determine the strength of inhibition evoked by PBSt group I afferents on the H-reflex, the amplitude of the conditioned H-reflex was expressed as a percentage of the amplitude of the unconditioned test reflex and was plotted against the C-T interval. To determine the strength of inhibition, the amplitude of the PBSt-evoked DRP was subtracted from the conditioned Tib-DRP to isolate the remaining Tib-DRP (Eccles *et al.*, 1963b; Wall & Lidierth, 1997). The amplitude of the conditioned Tib-DRP response expressed as a percentage of the unconditioned response was then averaged (n=8-10 traces) and plotted against the C-T interval.

## STATISTICS

One-way ANOVA followed by Holm-Sidak *post hoc* test (or Kruskal-Wallis followed by Dunn’s *post hoc* test if normality or equal variance test failed) was used to determine significant differences between groups unless stated below. To address the effect of the conditioning PBSt stimulation on Tib-DRP or H-reflex amplitude, a two-way repeated measures ANOVA followed by a Holm-Sidak *post hoc* test was performed to determine significant differences between groups, the C-T interval and their interaction. Results are presented as mean values ± standard deviation (SD). A linear regression analysis was used to correlate the conditioned H-reflex and DRP amplitude evoked by the same stimulation to the tibial nerve. Statistical analysis was performed using Sigma Plot software version 14.0 (SYSTAT software). For all statistical tests, the effects were considered significant when *P* < 0.05.

## RESULTS

### Chronic SCI decreases transmission in PAD pathways activated by PBSt afferents

To investigate the effect of chronic SCI and potential benefits of step-training on the strength of presynaptic inhibition, we first recorded dorsal root potentials (DRPs) evoked by the stimulation of 1) the posterior biceps-semitendinosus (PBSt) nerve, a knee flexor and hip extensor, known to be an efficient source of presynaptic inhibition (Eccles *et al.*, 1962c; Rudomin & Schmidt, 1999; Côté & Gossard, 2003), and 2) the tibial nerve, a mixed nerve containing both cutaneous and muscle afferents innervating the ankle and the plantar surface of the foot. DRPs were recorded from a cut L4 dorsal rootlet in response to a stimulation of the tibial nerve (Tib-DRP) or of the PBSt nerve (PBSt-DRP) (**Fig. 1*A***).

The properties of the recorded DRPs are depicted in Table 1. Chronic SCI did not significantly affect the latency to reach peak DRP (Tib-DRP: ***F*_2.18_ = 1.750; *P* = 0.202;** PBSt-DRP: ***F*_2.18_ = 2.145; *P* = 0.146**, one-way ANOVA). However, the stimulation intensity required to evoke a DRP of maximal amplitude (DRP_max_) was different between groups, but only when induced by a stimulation to PBSt (***H*_2.18_ = 13.986; *P* < 0.001,** Kruskal-Wallis) and not the tibial nerve (***F*_2.18_ = 0.106; *P* = 0.9,** one-way ANOVA). Overall, the stimulation intensity required to evoke PBST-DRP_max_ almost doubled after chronic SCI as compared to Controls from 2.32T ± 0.49 to 4.30T ± 0.55 (***P* < 0.05,** Dunn’s). This increase was independent from the stimulation intensity required to evoke an incoming afferent volley, as the CDP was not different between groups whether evoked by the tibial (***H*_2.18_ = 3.262; *P* = 0.196,** one-way ANOVA on Ranks) or the PBSt nerve (***F*_2.18_ = 0.686; *P* = 0.516,** one-way ANOVA).

**Table 1.** Properties of DRPs evoked by PBSt or tibial nerve stimulation.

|  |  | **PBST DRP<sub>max</sub>** |  |  | **Tibial DRP<sub>max</sub>** |  |  |
| --- | --- | --- | --- | --- | --- | --- | --- |
|  | n= | **Threshold**<br>**for CDP**<br>**(uA)** | **Stimulation**<br>**intensity**<br>**(xCDP)** | **Latency-to-**<br>**peak**<br>**(ms)** | **CDP**<br>**threshold**<br>**(uA)** | **Stimulation**<br>**intensity**<br>**(xCDP)** | **Latency-to-**<br>**peak**<br>**(ms)** |
| **Control** | 8 | 16.24 ± 8.72 | 2.32 ± 0.49 | 26.21 ± 2.16 | 7.13 ± 5.47 | 2.39 ± 0.41 | 20.86 ± 2.32 |
| **SCI** | 7 | 11.01 ± 6.17 | 4.30 ± 0.55*** | 23.89 ± 2.77 | 9.81 ± 2.48 | 2.34 ± 0.41 | 19.59 ± 1.91 |
| **SCI + Step-Training** | 6 | 15.00 ± 11.46 | 2.65 ± 0.55 | 23.61 ± 2.51 | 7.72 ± 4.10 | 2.29 ± 0.27 | 18.94 ± 1.43 |

Values are mean ± SD. DRP, Dorsal root potential; CDP, cord dorsum potential. \*\*\* P<0.001 (vs Control and SCI + Step-training).

The DRP represents the compound potential from activated afferents in the recorded rootlet at a given stimulation intensity. As the amplitude of the DRP reflects the amount of PAD evoked by the stimulated primary afferents (Lucas-Osma *et al.*, 2018), we therefore sought to investigate whether chronic SCI affects the recruitment gain of DRPs evoked by tibial and PBSt afferents. **Figure 2*A*** illustrates examples of DRPs recorded in response to stimulations of increasing intensity in a Control, SCI and a SCI + Step-training animal. The DRP amplitude augmented with stimulation intensity, whether evoked by PBSt or the tibial nerve, and the resulting inputoutput curve was fitted to a sigmoid function (**Fig. 2*B***). All animals displayed input-output relationships that tightly fit a sigmoid function (***P* < 0.001**) with individual R^2^ ranging from 0.93-0.98 in Controls, 0.94-0.99 in chronic SCI and 0.90-0.99 in chronic SCI + Step-training animals. Function parameters were then averaged by group (**Fig. 2*C-E***), and the sigmoid functions for each group plotted with a 95% confidence interval (**Fig. 2*F-G***). Overall, chronic SCI did not significantly affect DRP_max_ evoked by either the tibial (***F*_2,18_ = 0.642; *P* = 0.538**) or the PBSt nerve (***F*_2,18_ = 1.049; *P* = 0.371,** one-way ANOVA) whether the animals were step-trained or not (**Fig. 2*C***) However, the stimulation intensity required to reach 50% of DRP_max_ (s50) was significantly different between groups (**Fig. 2*D*)** both when evoked by the tibial (***F*_2,18_ = 7.457; *P* = 0.004**) or PBSt (***H*_2,18_ = 15.068; *P* < 0.001,** Kruskal-Wallis). Chronic SCI increased the s50 from 1.26T ± 0.08 to 1.39T ± 0.13 as compared to control when evoked by the tibial (***P* = 0.020,** Holm-Sidak) and from 1.35T ± 0.11 to 2.10T ± 0.38 when evoked by PBSt (***P* < 0.05,** Dunn’s). These results are further supported by a significant difference in the slope of the PBSt-DRP evoked input-output curve (***F*_2,18_ = 12.617; *P* < 0.001,** one-way ANOVA) between groups (**Fig. 2*E*)**, with chronic SCI significantly decreasing the steepness of the slope (***P* < 0.001,** Holm-Sidak). In contrast, the slope of Tib-DRPs input-output curve was not different between groups (**Fig. 2*E***; ***F*_2,18_ = 3.162; *P* = 0.067,** one-way ANOVA). Overall, our results suggest that although chronic SCI does not affect the amplitude of the maximal DRP that can be evoked, it decreases the amplitude of DRPs generated at submaximal stimulation intensity, specifically when evoked by PBSt (but not the tibial).

**Figure 2.**
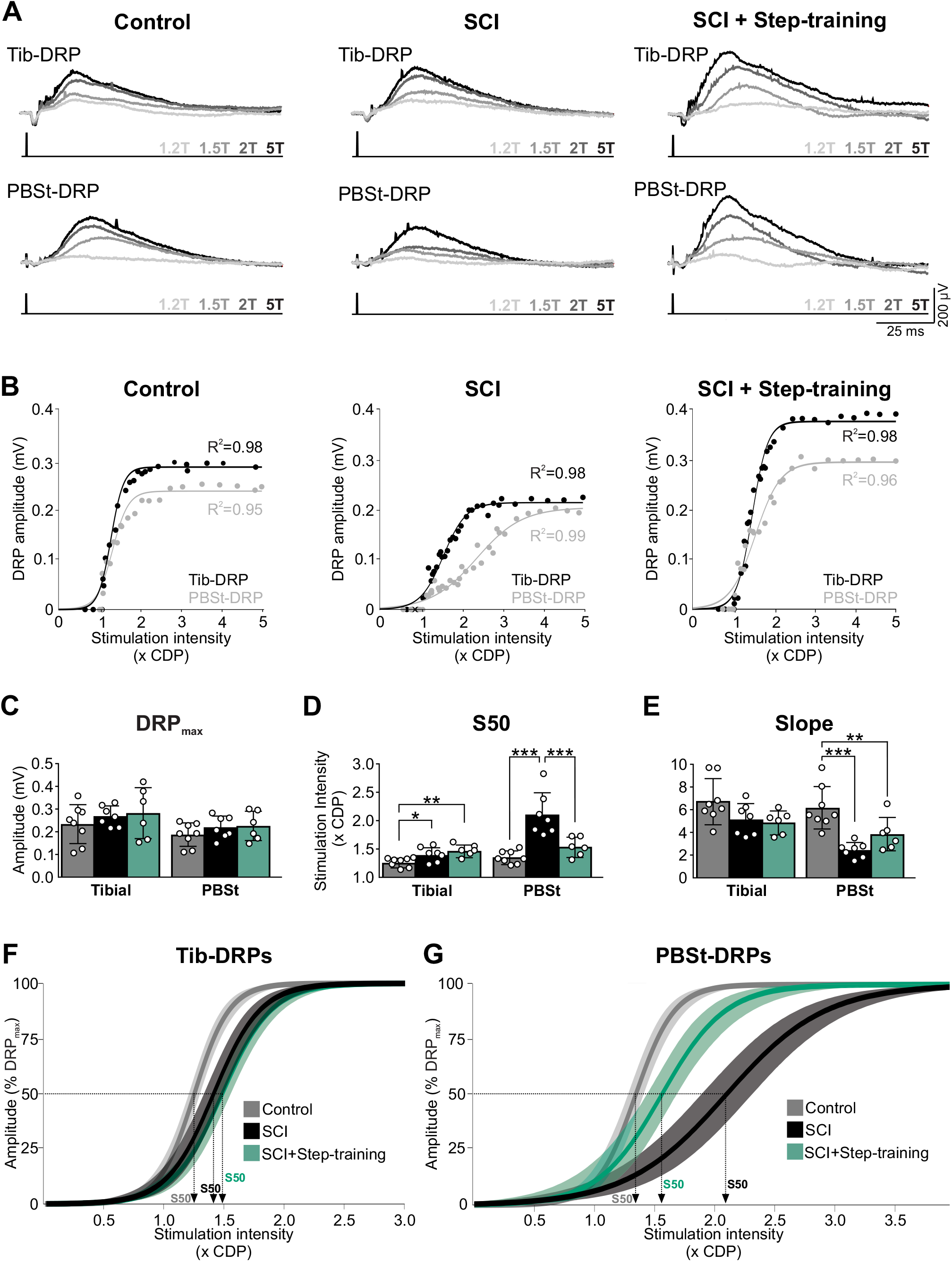
Chronic SCI and step-training specifically modulates DRPs evoked by PBSt group I muscle afferents. ***A*,** Example of DRPs evoked by the tibial (top) or PBSt (bottom) nerves at different stimulation intensities (1.2, 1.5, 2, or 5T) in a Control, a SCI and a SCI + Step-training animal. ***B*,** Example of the individual sigmoid function that was fitted to inputoutput relationships for each animal. Overall, all animals displayed curves that tightly fitted a sigmoid function (*P* < 0.001) with R^2^ ranging from 0.93-0.98 in Controls, 0.94-0.99 in SCI and 0.90-0.99 in SCI + Step-training. ***C-E*,** Chronic SCI did not significantly affect the amplitude of Tib-DRP_max_ (*P* = 0.538) and PBst-DRP_max_ (*P* = 0.371) whether the animals were step-trained or not (C). The stimulation intensity to reach 50% of DRP_max_ (s50) when evoked by the tibial (*P* = 0.004) or PBSt nerve (*P* < 0.001) was different across groups (D). Chronic SCI significantly increased s50 as compared to controls (tibial, ***P* = 0.020;** PBSt, ***P* < 0.001**). Step-training had no effect on s50 when DRPs were evoked by the tibial nerve (***P* = 0.225 vs. SCI**) but prevented the injury-dependent increase in s50 when evoked by PBSt (**p<0.001**) with values similar to Controls (***P* = 0.182**). The slope of PBST-DRPs input-output relationship was different across groups only when evoked by PBSt (***P* < 0.001**), not tibial nerve (***P* = 0.067**) (E). Chronic SCI decreased the slope (*P* < 0.001), while step-training had no further effect (***P* = 0.100 vs. SCI**). Results are expressed as mean ± SD, one-way ANOVA followed by Holm-Sidak *post hoc* test (Control, n=8; SCI, n=7; SCI + Step-training, n=6). CDP, cord dorsum potential; \**P* < 0.05; ** *P* < 0.01; \*\*\**P* < 0.001. ***F-G*,** Sigmoid function resulting from group averages is illustrated with 95% confidence interval for Tib- and PBSt-DRPs.

### Step-training restores the transmission in PAD pathways activated by PBSt afferents after chronic SCI

Among the beneficial effect of step-training after SCI is a decrease in hyperreflexia and spasticity and improvement in spinal reflex modulation that is believed to rely, at least in part, to a restoration in presynaptic inhibition (Bouyer & Rossignol, 2003; Côté & Gossard, 2004). We sought to investigate whether step-training more specifically prevented the detrimental effect of chronic SCI on Tib- and PBSt-DRP amplitude. As reported in the previous paragraph, no group difference was observed in latency to reach peak DRP amplitude, nor in the stimulation intensity required to reach DRP_max_, suggesting that step-training did not affect those properties (**Table 1**). Also, step-training did not affect any of the parameters of the input-output relationship of evoked Tib-DRPs (**Fig. 2*C-E***). However, the stimulation intensity required to evoke PBSt-DRP_max_ was decreased after step-training as compared to non-exercised SCI animals from 4.30T ± 0.55 to 2.65T ± 0.55 **(*P* < 0.05**, Dunn’s), and was no different from Controls (**P > 0.05,** Dunn’s) (**Table 1**). Step-training restored the stimulation intensity required to reach 50% of DRP_max_ (**Fig. 2*D, P* < 0.001**) with values similar to control (***P* = 0.182,** Dunn’s). These results indicate that step-training contributes to preventing the decrease in DRP amplitude originating from PBSt afferents after chronic SCI.

### The disruption in PAD pathways activated by PBSt group I muscle afferents is specifically affected by chronic SCI

The decrease in s50 after chronic SCI in the absence of a decrease in the maximal DRP amplitude evoked by PBSt suggests a decrease in DRP amplitude at submaximal intensity, in the range of large PBSt group I muscle afferent activation. Previous studies have suggested that stimulation intensities < 2T and frequencies >200Hz specifically increase the activation of PAD pathways by group I muscle afferents (Tasaki, 1953; Eccles *et al.*, 1963b). To specifically address the role of PAD pathways activated by group I muscle afferents, without engaging group II afferents, we measured the amplitude of the DRP evoked in L4 dorsal rootlet by a tetanic stimulation to PBSt nerve at submaximal group I afferents strength (4p, 250Hz, 1.5T). **Figure 3*A*** shows an example of DRPs evoked by PBSt group I afferents in a Control and chronic SCI animals with or without step-training, with the associated DRP_max_ illustrated for reference. We found a significant difference in the relative DRP amplitude between groups **(*F*_2.18_ = 14.336; P < 0.001,** one-way ANOVA). Chronic SCI decreased the amplitude of DRPs evoked by PBSt group I afferents (***P* = 0.002,** Holm-Sidak). In addition, step-trained animals displayed significantly larger PBSt-DRPs amplitude than chronic SCI (***P* < 0.001**), and no significant difference with control animals **(*P* = 0.058,** Holm-Sidak), suggesting a return of the presynaptic inhibition originating from PBSt group I afferents with step-training (**Fig. 3*B***). Because the size of the DRP is highly sensitive to the number of afferents in the recorded rootlet and the distance between the recording electrode and the cord, we chose to express the amplitude of the DRP evoked by PBSt group I afferents as a function of the DRP_max_ recorded in the same experimental conditions. It is important to note that a significant difference in DRP *absolute* amplitude (in mV, *not shown*) was also observed between groups **(*F*_2,18_ = 16.315; P < 0.001,** one-way ANOVA) with a similar decrease in SCI animals (***P* = 0.026,** Holm-Sidak) and restoration of amplitude after step-training, with DRPs significantly larger than chronic SCI (***P* < 0.001,** Holm-Sidak) and also larger than control animals **(*P* = 0.002,** Holm-Sidak).

**Figure 3.**
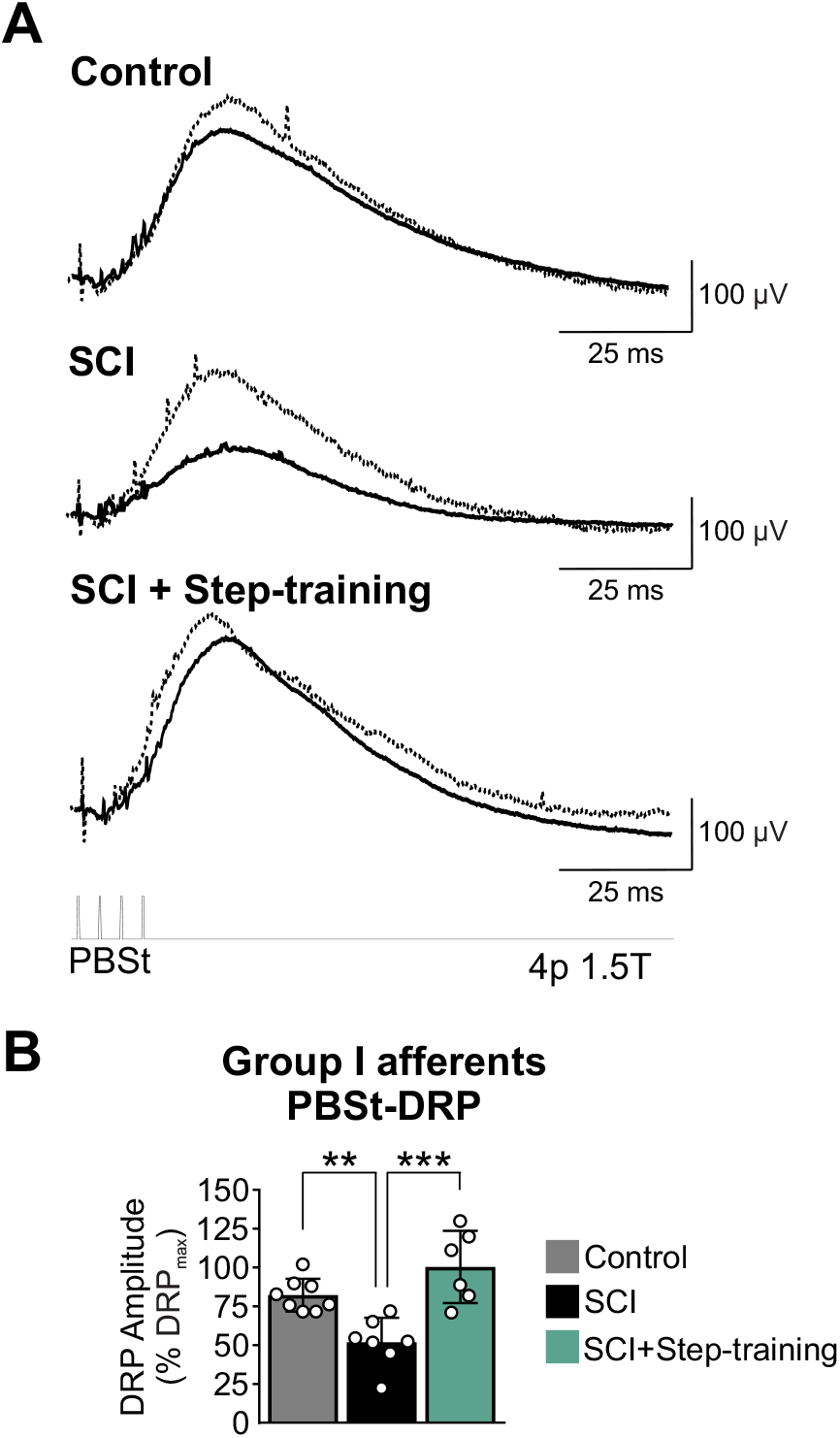
Disruption is the transmission of group I PBSt afferents in PAD pathways after chronic SCI is improved by step-training. ***A*,** Representative traces of DRPs evoked by a stimulation to PBSt nerve at group I afferent strength (4p, 1.5T, 250Hz) in a Control, a SCI and a SCI + Step-training animal. For reference purpose, the dotted line illustrates DRP_max_ in the same animal. ***B*,** The amplitude of DRPs evoked by PBSt group I afferents was different across groups **(*P* < 0.001).** Chronic SCI decreased DRP amplitude (*P* = 0.002). This decrease was prevented by step-training with values similar to Controls (*P* = 0.058) but significantly larger than chronic SCI animals (***P* < 0.001)**. Results are expressed as mean ± SD, one-way ANOVA followed by Holm-Sidak *post hoc* test ** *P* < 0.01; *** *P* < 0.001.

### Step-training promotes the inhibition of Tib-DRPs induced by PBSt group I afferents after chronic SCI

To assess the strength and time course of presynaptic inhibition originating from PBSt group I afferents, we used a condition-test protocol and recorded DRPs evoked by a stimulation to the tibial nerve when preceded or not by a conditioning stimulation of PBSt group I afferents (Eccles *et al.*, 1962b; 1963b) (**Fig 4*A*)**. **Figure 4*B*** illustrates that after chronic SCI, there is minimal effect of a conditioning stimulation to PBSt, unless the animal was step-trained. Overall, there was a significant difference between groups (***F*_2,171_ = 9.985; *P* = 0.001**), C-T intervals (***F*_9,171_ = 58.299; *P* < 0.001**), and an interaction between groups and C-T intervals (***F*_18,171_ = 1.847; *P* = 0.023,** two-way RM ANOVA). The inhibition of the Tib-DRP exerted by PBSt group I afferents was significantly decreased after chronic SCI animals at C-T intervals ranging from 10 to 60ms as compared to Controls (**Fig. 4*C, P* < 0.05,** Holm-Sidak). Step-training restored inhibition at intervals ranging from 10 to 50ms (***P* < 0.05,** Holm-Sidak) and was similar to Control (***P* > 0.05,** Holm-Sidak). These results suggest that presynaptic inhibition induced by PBSt group I afferents is decreased after chronic SCI unless the animals are step-trained indicating that activity-based therapies may contribute to restore inhibition in this pathway.

**Figure 4.**
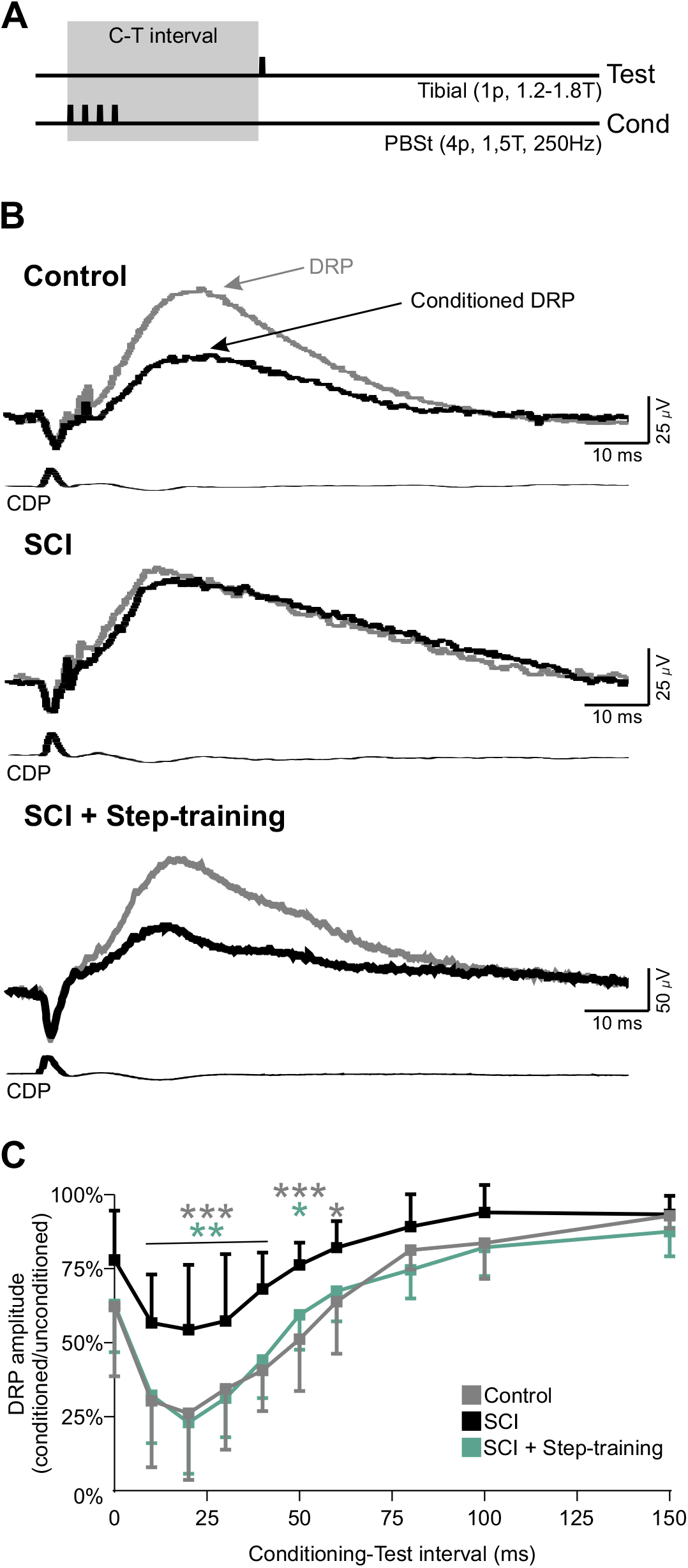
Step-training improves the inhibition induced by PBSt group I afferents on L4-DRPs evoked by the tibial nerve after chronic SCI. **A)** Conditioning-test protocol used to estimate the level of inhibition evoked by a conditioning stimulation to PBSt group I afferents (4p, 250Hz, 1.5T) on Tib-DRPs (1p, 0.2 Hz, 1.2-1.8T). **B,** DRPs evoked by the tibial nerve when preceded (black) or not (gray) by a conditioning stimulation to PBSt (C-T: 50 ms) in a Control, chronic SCI and SCI + Step-training animal. The inhibitory effect of the conditioning stimulation is not present after chronic SCI. **C,** Overall, chronic SCI significantly decreases the strength of inhibition at C-T intervals ranging from 10 to 60ms as compared to Controls (***P* < 0.05,** gray stars). This decrease was not observed in step-trained animals with values similar to Control and inhibition significantly larger than SCI animals at intervals ranging from 10 to 50ms (***P* < 0.05**, green stars). Two-way RM ANOVA followed by Holm-Sidak *post hoc* test, *p<0.05; **p<0.01; ***p<0.001; Control (n=8), SCI (n=7), SCI + Step-training (n=6). C-T: Conditioning-Test.

### Presynaptic inhibition exerted by PBSt group I muscle afferents contributes to plantar H-reflex hyperexcitability after chronic SCI

In order to investigate if the decrease in presynaptic inhibition contributes to hyperreflexia, we assessed the effect of a conditioning stimulation to PBSt on DRPs and H-reflexes evoked by a stimulation to the tibial nerve simultaneously. We first examined the excitability of the H-reflex by analyzing the general features of the M-waves and H-reflexes. Chronic SCI did not alter the latency, the threshold for activation, nor the maximal amplitude of the M-wave and H-reflex (**Table 2**). **Figure 5*A*** illustrates examples of M-wave and H-reflex recruitment curves in animals from the Control, chronic SCI and chronic SCI + step-training groups. The recruitment curves of the M-wave and H-reflex were individually fitted to a sigmoid function for each animal. All animals displayed recruitment curves that tightly fitted a sigmoid function (***P* < 0.001**), with R^2^ ranging from 0.93-0.99 for Control, 0.90-0.99 for SCI and 0.91-0.99 for the SCI + Step-training group. Function parameters were then averaged by group and the resulting sigmoid functions for each group plotted with a 95% confidence interval (**Fig. 5*B-C***). Further analysis of parameters of the sigmoid function illustrate that chronic SCI did not change the maximal H-reflex amplitude (**Fig. 5*D***; ***F*_2.18_ = 0.826; *P* = 0.454,** one-way ANOVA) nor the stimulation intensity to obtain 50% of Hmax (**Fig. 5*E*; F_2.18_ = 0.510; *P* = 0.609,** one-way ANOVA) whether the animals were step-trained or not. However, the slope of the H-reflex sigmoid function was significantly different between groups (**Fig. 5*F*; *F*_2.18_ = 7.682; *P* = 0.004**, one-way ANOVA). Chronic SCI dramatically increased the slope of the H-reflex recruitment curve as compared to Controls (***P* = 0.002**) and Step-trained animals (***P* = 0.015,** Holm-Sidak). This observation is unlikely due to changes at the neuromuscular junction as we found no significant difference in Mmax (**Fig. 5*D*; *F*_2.18_ = 0.534; *P* = 0.595,** one-way ANOVA), s50 (**Fig. 5*E*; *F*_2.18_ = 0.951; *P* = 0.405,** one-way ANOVA) or the slope of the M-wave recruitment curve (**Fig. 5*F*; *F*_2.18_ = 2.463; *P* = 0.113,** oneway ANOVA). Altogether, these results suggest an increase in the recruitment gain of the motor pool by tibial afferents after chronic SCI that is normalized by step-training.

**Figure 5.**
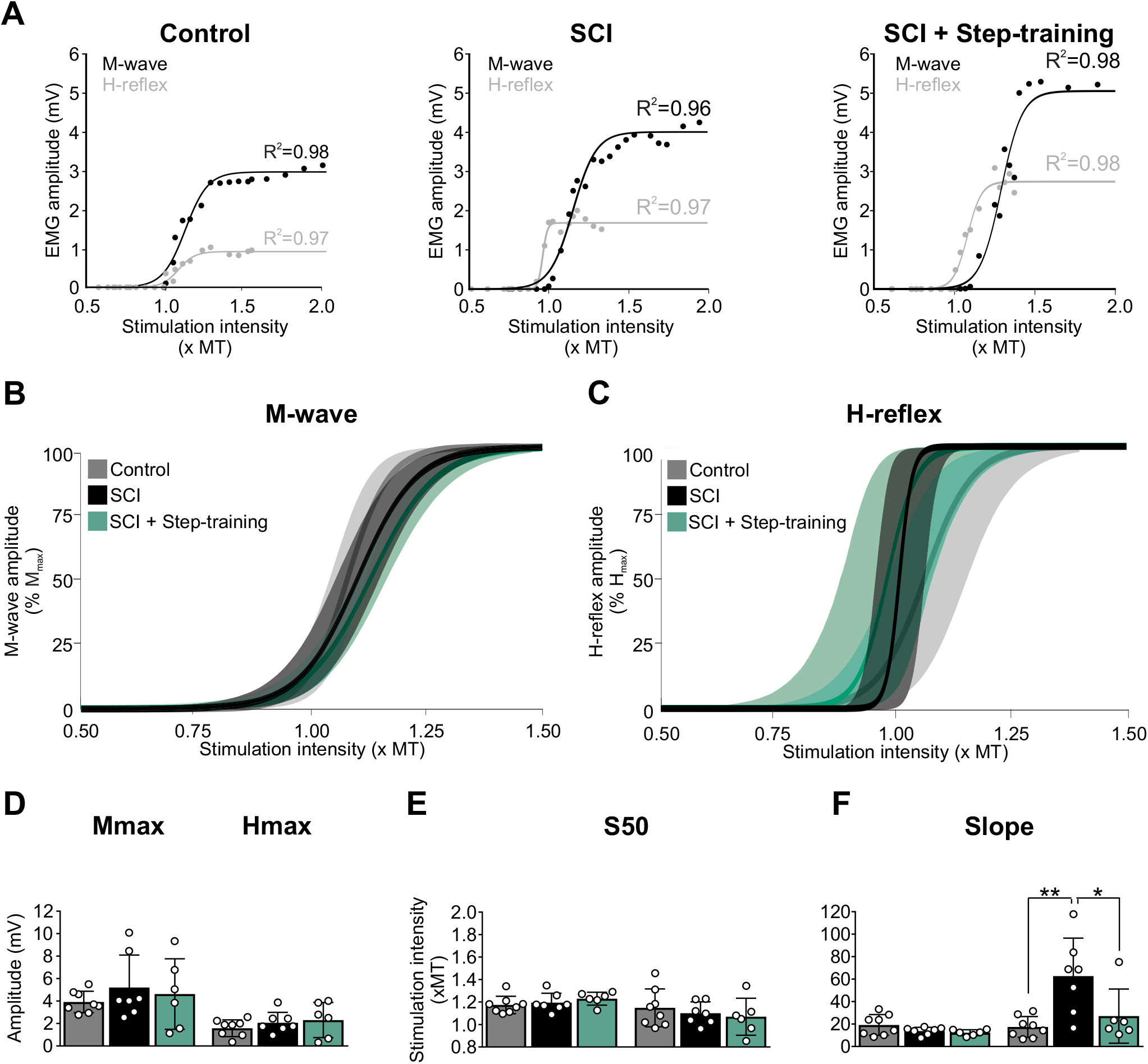
H-reflex excitability is impaired following chronic SCI and normalized by step-training. **A)** Example of M-wave and H-reflex recruitment curves in a Control, a SCI and a SCI + Step-training animal. Sigmoid functions were individually fitted to the recruitment curves. Overall, all animals displayed curves that tightly fitted a sigmoid function (***P* < 0.001**) with R^2^ ranging from 0.93-0.99 in Controls, 0.90-0.99 for SCI and 0.91-0.99 for the SCI + Step-training animals. **B-C)** The sigmoid function resulting from group averages is illustrated with 95% confidence interval for both the M-wave (**B**) and the H-reflex (**C**). **D-F)** Chronic SCI did not alter the Hmax and Mmax amplitude (D) (***P* = 0.454 and *P* = 0.595** respectively) and the stimulation intensity to reach 50% of the maximal amplitude (E) (***P* = 0.609 and *P* = 0.405** respectively) whether the animals were step-trained or not. However, the slope of the H-reflex sigmoid function was significantly different across groups (F) (***P* = 0.004**). Chronic SCI increased the steepness of the slope of the recruitment curve for the H-reflex as compared to Controls (***P* = 0.002**) and step-trained animals (***P* = 0.015**), while step-training preserved a slope similar to Controls (***P* = 0.465**). The slope of the M-wave recruitment curve was similar across groups (***P* = 0.113).** One-way ANOVA followed by Holm-Sidak *post hoc* test (Control n=8; SCI n=7; SCI + Step n=6). MT, motor threshold; *p<0.05; **p<0.01.

**Table 2.** Properties of the M-Wave and H-Reflex.

|  |  | **M-wave** |  |  | **H-reflex** |  |  |
| --- | --- | --- | --- | --- | --- | --- | --- |
|  | n= | **Motor threshold**<br>**(uA)** | **Latency**<br>**(ms)** | **M<sub>max</sub>**<br>**(xMT)** | **Threshold**<br>**(x MT)** | **Latency**<br>**(ms)** | **H<sub>max</sub>**<br>**(xMT)** |
| **Control** | 8 | 10.49 ± 9.23 | 1.88 ± 0.33 | 1.46 ± 0.26 | 0.96 ± 0.12 | 8.20 ± 0.79 | 1.45 ± 0.24 |
| **SCI** | 7 | 12.21 ± 2.87 | 2.11 ± 0.16 | 1.48 ± 0.21 | 1.03 ± 0.13 | 8.00 ± 0.38 | 1.21 ± 0.14 |
| **SCI + Step-**<br>**Training** | 6 | 10.43 ± 5.84 | 2.02 ± 0.49 | 1.52 ± 0.27 | 0.94 ± 0.17 | 7.93 ± 0.59 | 1.34 ± 0.22 |

Values are mean ± SD. MT, motor threshold.

**Figure 6*A*** illustrate an example of conditioned and unconditioned H-reflexes from a Control, a SCI and a SCI + step-training animal at 10 and 50ms C-T intervals. The conditioning stimulation to PBSt dramatically decreases the H-reflex amplitude at both C-T intervals in the Control and SCI + Step-training animals, but not in the chronic SCI. Overall, there was a significant difference in the inhibition of the H-reflex between groups (***F*_2.171_ = 32.056; *P* < 0.001**), C-T interval (***F*_9.171_ = 29.865; *P* < 0.001**), and an interaction between groups and C-T intervals (***F*_18.171_ = 2.506; *P* = 0.001,** two-way RM ANOVA). For clarity, these significant differences across C-T intervals within a given group are not illustrated in **Figure 6B**. The lack of modulation after chronic SCI is further supported by a decrease in H-reflex inhibition between chronic SCI as compared to Controls (***P* < 0.01,** grey stars) and step-trained animals (***P* < 0.05,** green stars) at intervals ranging from 10-150ms. Hence, step-training promoted H-reflex inhibition after chronic SCI for intervals ranging from 10-150ms to values similar to Controls, except for the 20-30ms intervals where step-trained animals displayed more inhibition than SCI, but less than Controls (***P* < 0.05**). Because the quantification of H-reflex inhibition by the conditioning stimulation is critically dependent on the amplitude of the test H-reflex response (Crone *et al.*, 1990), we further insure that the amplitude of the unconditioned H-reflex was similar across groups (**Fig. 6*C*, *F*_2,18_ = 2.022; *P* = 0.161,** one-way ANOVA).

**Figure 6.**
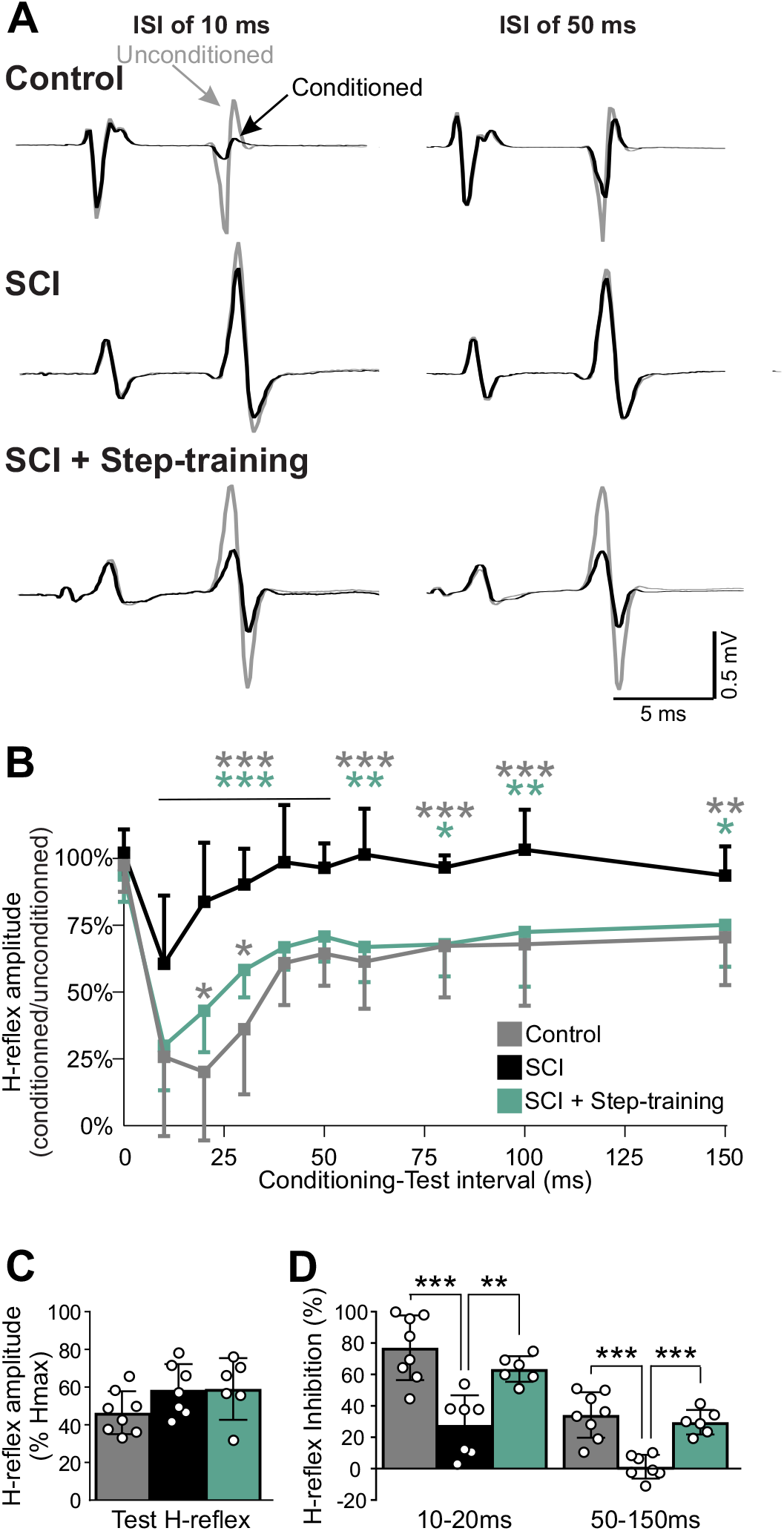
Step-training promotes H-reflex inhibition after chronic SCI. **A)** Representative recordings of H-reflexes evoked by a stimulation to the tibial nerve with or without a conditioning stimulation to PBSt in a Control, SCI and SCI + Step-training animal at C-T intervals of 10ms and 50ms. **B)** There was a significant difference in the strength of inhibition of the H-reflex between groups (**F_2,171_=32.056; p<0.001**), C-T interval (**F_9,171_=29.865; *P* < 0.001**), and an interaction between groups and C-T intervals (**F_18,171_=2.506; *P* = 0.001**). For clarity purpose, only significance between groups, but not intragroup is illustrated. Chronic SCI significantly decreased the depression of the H-reflex as compared to Control at intervals ranging from 10-150ms (***P* < 0.01**). In addition, the strength of inhibition in step-trained animals was larger than in non-trained animals (***P* < 0.05,** green stars) and similar to Controls at similar C-T intervals. Two-way RM ANOVA followed by Holm-Sidak *post hoc* test. C-T, Conditioning-Test. **C)** The amplitude of the unconditioned H-reflex was similar between experimental groups (***P* = 0.395**). One-way ANOVA. **D)** There was a significant decrease in postsynaptic (C-T intervals 10-20ms; **F_2,18_ = 15.567, *P* < 0.001**) and presynaptic inhibition (C-T intervals 50-150ms; **F_2,18_ = 16.597, *P* < 0.001**). Chronic SCI decreased the strength of postsynaptic (***P* < 0.001,** gray stars) and presynaptic inhibition (***P* < 0.001,** gray stars), while step-training promoted their recovery (respectively ***P* = 0.002** and ***P* < 0.001,** green stars). One-way ANOVA followed by Holm-Sidak *post hoc* test. \**P* < 0.05; ** *P* < 0.01; *** *P* < 0.001.

PBSt afferents simultaneously activate GABAergic pathways involved in PAD generation, but also contribute to postsynaptic inhibitory pathways leading to IPSPs in motoneurons (Hultborn *et al.*, 1987a; Stuart & Redman, 1992; Pierce & Mendell, 1993; Hughes *et al.*, 2005), both of which contribute to modulating the H-reflex amplitude. We therefore sought to isolate presynaptic inhibitory effects from postsynaptic inhibition. To do so, we included the longest C-T intervals (50-150ms) to isolate presynaptic inhibition, while postsynaptic inhibition was estimated at shorter intervals (10-20ms) (Eccles *et al.*, 1961; 1962c). There was a significant difference in postsynaptic (***F*_2,18_ = 15.567; *P* < 0.001,** one-way ANOVA) and presynaptic inhibition (***F*_2,18_ = 16.597; *P* < 0.001,** one-way ANOVA). Chronic SCI decreased postsynaptic (***P* < 0.001**) and presynaptic inhibition (***P* < 0.001**) while step-training promoted their recovery (respectively ***P* = 0.002** and ***P* < 0.001,** Holm-Sidak) (**Fig. 6*D***). These results suggest that chronic SCI reduces inhibition originating from group I PBSt afferents onto tibial Ia afferents projecting to motoneurons (presynaptic inhibition) and onto interosseous motoneurons (postsynaptic inhibition). Step-training counteracted the effects by promoting the recovery of both the postsynaptic and presynaptic inhibition.

### Chronic SCI disrupts the association between DRP and H-reflex inhibition

The inhibition of the monosynaptic reflex and Tib-DRP evoked by PBSt group I muscle afferents are believed to rely on similar mechanisms due to the similarity in the strength and duration of their inhibition (Eccles *et al.*, 1962b-c; 1963b). We further performed a regression analysis to assess if the strength of inhibition evoked by PBSt group I afferents on Tib-DRPs is predictive of the strength of inhibition incurred on the plantar H-reflex (**Fig. 7**). The inhibition of the conditioned Tib-DRP at a given C-T interval significantly co-varied with the inhibition of the conditioned H-reflex in Controls (R^2^: 0.244, ***P* < 0.001**) and SCI + Step-training (R^2^: 0.264, ***P* < 0.001,** linear regression), but not in chronic SCI animals (R^2^: 0.0150, ***P* = 0.301**). The lack of correlation in SCI animals is further supported by the clustering of data in the top right corner with little inhibition of both the DRP and H-reflex amplitude in chronic SCI animals (***Fig. 7B***) contrary to Controls (Fig. ***7A***) and Step-trained animals (***Fig. 7C***). Together, these results suggest that the inhibition evoked by PBSt group I afferents on Tib-DRP is predictive of the inhibition incurred on the plantar H-reflex under Control conditions but is lost after chronic SCI unless the animals are step-trained.

**Figure 7.**
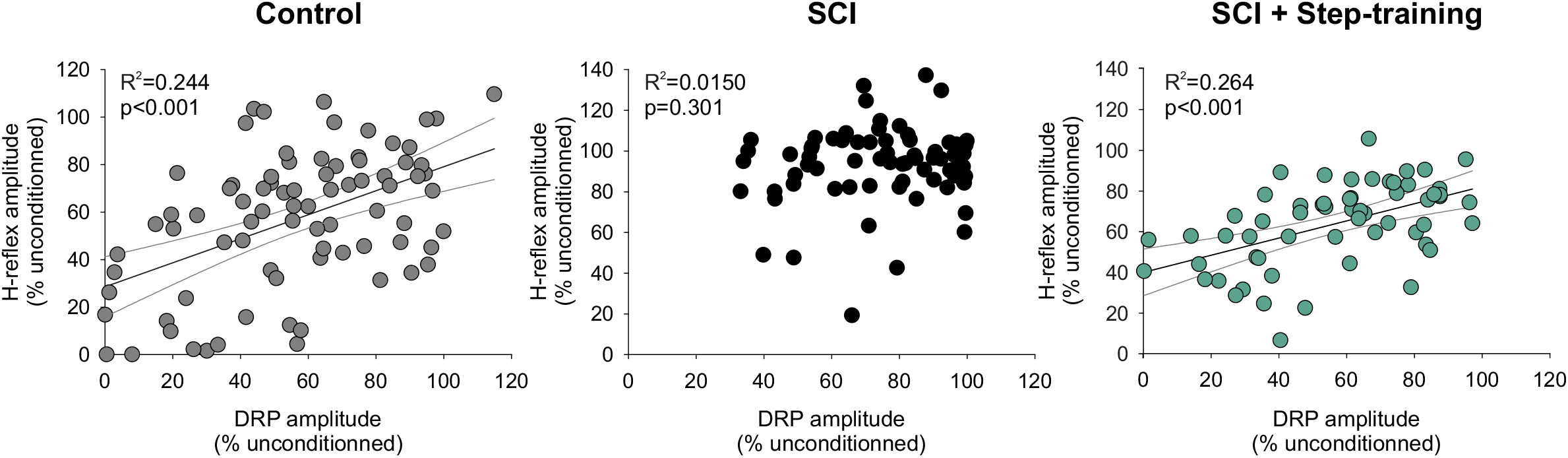
Chronic SCI disrupts the relationship between DRP amplitude and H-reflex inhibition unless the animal is step-trained. The linear relationship between the amplitude of the H-reflex and DRP conditioned by a stimulation to PBSt in Control animals (R^2^: 0.244, ***P* < 0.001**) is not present in chronic SCI animals (R^2^: 0.0150, ***P* = 0.301**), but restored by step-training (R^2^: 0.264, ***P* < 0.001**). Regression Analysis.

## DISCUSSION

In rodents, most studies investigating presynaptic inhibition derive from reduced preparations of adult spinal cords *ex vivo* (Lucas-Osma *et al.*, 2018), neonates in which the development of the nervous system is not yet fully complete (Hochman *et al.*, 2010; Garcia-Ramirez *et al.*, 2014), or adult anaesthetized preparation (Wall & Lidierth, 1997). Here, we show a decrease in the activation of lumbar PAD pathways associated with a reduction in presynaptic inhibition of the H-reflex *in vivo* in the adult decerebrated rat after a chronic low thoracic transection. More precisely, we have demonstrated that the activation of flexor muscle group I PAD pathways activated by PBSt group afferents is significantly disrupted after chronic SCI and contributes to decrease presynaptic inhibition of the plantar H-reflex in association with an increased recruitment of motor pools. We further show that the repetitive activation of sensory afferents provided by step-training reestablishes presynaptic inhibition originating from PBSt group I afferents on the H-reflex through restoring transmission in PAD pathways after chronic SCI.

### Afferent control of presynaptic inhibition from PBSt group I afferents is selectively impaired after chronic SCI

Pathways mediating presynaptic inhibition provide a powerful mechanism by which the incoming sensory information flooding the spinal cord is regulated before it even reaches the first central synapse (reviewed in Rudomin & Schmidt, 1999; Rudomin, 2009). Presynaptic inhibition is mainly produced via primary afferent depolarization (PAD) of intraspinal terminals, which can be measured experimentally following the antidromic electrotonic spread to the dorsal roots and recorded as a dorsal root potential (DRP) (Barron & Matthews, 1938; Eccles *et al.*, 1963b). Presynaptic inhibition is under the control of both supraspinal centers and primary afferents in order to gate afferent feedback and to adjust reflex excitability to meet task requirements (Rudomin & Schmidt, 1999). While the supraspinal control of PAD pathways has been well described, little is known about maladaptive plasticity triggered by chronic SCI when supraspinal control is disrupted, and how it impacts sensory-evoked presynaptic inhibitory mechanisms.

In our experiment, chronic SCI decreased the amplitude of DRPs evoked by tibial afferents to a much lesser extent than DRPs evoked by PBSt. Because the tonic descending inhibition of PAD pathways is acutely released by a transection in our Control decerebrated animals (Carpenter *et al.*, 1963; Quevedo *et al.*, 1993), the changes in DRP amplitude likely reflects changes in the transmission of released sensory PAD pathways. The highly specific nature of PAD pathway regulation suggests that primary afferents receive specific patterns of presynaptic inhibition with different terminals being selectively regulated (Lomeli *et al.*, 1998). The tibial nerve is a mixed nerve, innervating multiple muscles at the ankle joint while its distal branch innervates the plantar surface of the foot and contains mostly cutaneous afferents. Although the specific contribution of cutaneous and low threshold muscle afferents to Tib-DRP amplitude cannot be determined with certainty, the location of our electrode on the distal part of the tibial nerve suggests a contribution of mostly cutaneous origin (Eccles *et al.*, 1962b; Loeb, 1993; Frigon *et al.*, 2012). PAD pathways activated by cutaneous afferents primarily generate PAD on other cutaneous and Ib-II afferents (Janig *et al.*, 1968b, a; Harrison & Jankowska, 1989; Quevedo *et al.*, 1995; Enriquez *et al.*, 1996b, a) and receive tonic inhibition from supraspinal centers (Carpenter *et al.*, 1963; Quevedo *et al.*, 1993). Our results indicate that the DRPs evoked by tibial afferents are barely altered by chronic SCI and suggest that the increased gain of sensory afferents vs. supraspinal input on spinal networks after chronic SCI (Edgerton *et al.*, 2001) does not significantly alter the activation of tibial PAD pathways. In contrast, DRPs evoked by PBSt afferents were significantly smaller, suggesting that PAD pathways activated by hip extensor and/or knee flexor afferents are critically disrupted after chronic SCI. PBSt is a purely muscle nerve known to be an efficient source of presynaptic inhibition (Frank & Fuortes, 1957; Eccles *et al.*, 1962c; Côté & Gossard, 2003). While chronic SCI did not decrease the amplitude of PBSt-DRP_max_, the input-output curve was significantly shifted toward higher stimulation intensities, i.e. higher stimulation intensities were required to evoke PBSt-DRP_max_ and to reach s50 (***Fig. 2 and Table 1***). In decerebrated acutely transected cats, the recruitment of group Ia PBSt afferents is initiated at 1T and reaches a maximum ~1.5T. Higher intensity of stimulation is required to activate group Ib afferents (~1.4T), reaching maximal activation ~2T, which overlaps with the lowest threshold for group II afferents activation (~1.6T) (Eccles & Lundberg, 1959; Carpenter *et al.*, 1963). While we have not directly investigated the type of muscle afferents recruited, the stimulation intensity required to reach PBST-DRP_max_ was consistent with group I strength in Control animals and with group II afferents following chronic SCI (***Fig. 2G***).

Because the alteration in the amplitude of PBSt-evoked DRP is mainly observed at group I strength, but not seen when group II afferents were engaged, we stimulated PBSt at submaximal group I strength (1.5T) to minimize the contribution of group II afferents. Hence, stimulation at maximal group I strength (2T) or higher was shown to not only maximally activate group I PAD pathways, but also engage low threshold group II PAD pathways, which brings DRP amplitude closer to DRP_max_ values (Eccles & Lundberg, 1959; Naftchi *et al.*, 1979). Consistent with this, the additional recruitment of PBSt group II afferents barely increases the DRP amplitude, as group I afferents already evoked a large DRP that is close to maximal amplitude. In contrast, when DRPs evoked by group I afferents are small, the additional recruitment of group II afferents produces a large increase in DRP amplitude (Eccles *et al.*, 1961). This is likely due to the activation of separate pathways by PBSt group I and II afferents that respectively evoke PAD on Ia, Ib and cutaneous afferents or on group II afferents (Riddell *et al.*, 1993). Accordingly, our results suggest that PAD evoked by PBSt group I afferents on Ia, Ib and cutaneous afferents is specifically decreased after chronic SCI and that engaging PBSt group II afferents does not reveal any further change. Together, our results indicate that presynaptic inhibition evoked by PBSt group I muscle afferents is specifically altered after chronic SCI, while presynaptic inhibition evoked by PBSt group II or tibial cutaneous afferents (historically termed flexor reflex afferents) is unlikely to contribute. Whether decreased transmission in group I PAD pathway is the consequence of a change in interneuronal excitability, primary afferents or other factors remains to be determined.

### H-reflex hyperexcitability is associated with a concurrent increase in the recruitment of motor pools and impairment in presynaptic and postsynaptic inhibition after chronic SCI

The DRP represents the sum of PADs from afferents present in the recorded dorsal rootlet. However, unlike the PAD, DRPs do not yield information on the specific type of afferents receiving presynaptic inhibition. A convenient method of testing this is to study the effect of a conditioning stimulation known to evoke PAD on primary afferents in a specific pathway. The H-reflex, the electrical analogue of the stretch reflex, is commonly used to assess the excitability of the monosynaptic reflex. Because of its monosynaptic nature, the H-reflex prevents the possible involvement of a change in excitability of interneurons present in the reflex pathway under different conditions. The modulation of the H-reflex has been extensively investigated in the rat using repeated homonymous sensory activation (Thompson *et al.*, 1992; Côté *et al.*, 2014; Caron *et al.*, 2016), which includes presynaptic, homosynaptic and postsynaptic inhibitory mechanisms (Hultborn *et al.*, 1996; Zucker & Regehr, 2002).

Presynaptic inhibition of hindlimb muscle group Ia afferents is mainly under the influence of flexor muscle group I afferents (Eccles *et al.*, 1961; 1962a, c; 1963a-b). Last-order inhibitory interneurons involved in presynaptic inhibition of the monosynaptic reflex synapse onto terminals of Ia muscle spindle afferents (Hughes *et al.*, 2005) and are mostly activated by flexor muscle group I afferents, inhibited by flexor reflex afferents, and controlled by descending tracts (Jankowska, 2001). Presynaptic inhibition exerted by PBSt group I afferents on the triceps surae monosynaptic reflex in the cat includes a short-lasting inhibition with an onset around 5ms and reaching maximal amplitude around 30ms, followed by long-lasting inhibition which persists for over 100ms (Eccles *et al.*, 1962a). Intracellular motoneuron recordings, following the same stimulation, similarly showed that PBSt group I afferents induce 40ms IPSPs in triceps surae motoneurons (postsynaptic inhibition) and decreased monosynaptic EPSPs (presynaptic inhibition) (Frank & Fuortes, 1957; Hultborn *et al.*, 1987a; Stuart & Redman, 1992). In this study, the heteronymous inhibition of the plantar H-reflex by PBSt eliminates the possibility for post-activation depression, so that long C-T intervals (50-150ms) reflect presynaptic inhibition exerted on tibial Ia afferents and short C-T intervals (10-20ms) mainly reflect postsynaptic inhibition exerted on motoneurons, with a minimal contribution of presynaptic inhibition onto tibial Ia afferents. Our data show that chronic SCI did not modify the time course, but noticeably decreased the strength of both postsynaptic and presynaptic inhibition. Chronic SCI noticeably decreased the strength of both postsynaptic and presynaptic inhibition of the H-reflex. However, changes in motoneuronal properties can also influence the strength of H-reflex, which would not be reflected by measures of either pre- or postsynaptic inhibition (Kernell & Hultborn, 1990). An increase in the slope of the H-reflex recruitment curve is associated with hyperreflexia and routinely used as a clinical evaluation of spasticity (Biering-Sorensen *et al.*, 2006). Accordingly,our data show that chronic SCI increased the slope of the plantar H-reflex recruitment curve, suggesting a greater recruitment of motor pools by Ia afferents (Capaday, 1997). Under normal conditions, Henneman’s size principle suggest that Ia afferents first activate slow motor units, such that motor units producing the smallest force are recruited the earliest (Henneman, 1957; Burke & Rymer, 1976; Harrison & Taylor, 1981). However, increasing the recruitment gain of the motor pool induces a fast and synchronous activation of the entire motoneuronal pool (Nielsen *et al.*, 2019), which contributes to further increase the excitability of the H-reflex (Kernell & Hultborn, 1990). This effect can be the consequence of several factors, including, 1) homogenization of motor unit properties, 2) increased inhibition biased towards slow motor units, 3) increased excitation biased towards large faster motor units, 4) a combination of aforementioned factors (Kernell & Hultborn, 1990).

After SCI and other conditions inducing immobilization, the intrinsic properties of slow motor units change towards faster motor unit properties (Burnham *et al.*, 1997; Talmadge, 2000; Beaumont *et al.*, 2004; Cormery *et al.*, 2005), which would likely contribute to increase the recruitment gain of the motor pool and influence the strength of H-reflex inhibition (Kernell & Hultborn, 1990). To account for this possibility, we have simultaneously recorded the inhibition exerted by PBSt group I afferent onto the plantar H-reflex, which includes the motoneuronal response, and onto the Tib-DRP, which does not involve the motoneuronal response. We show that the inhibition exerted by PBSt group I afferents on both the H-reflex and Tib-DRP is decreased after chronic SCI, confirming that chronic SCI also decreases inhibition independently of motoneuron activation. In addition, the correlation between the strength of inhibition exerted by PBSt on Tib-DRP and on the H-reflex observed in Controls, was not present in chronic SCI animals. This lack of correlation further supports a disruption in presynaptic inhibition after SCI.

Together, our results suggest that flexor muscle group I afferents PAD pathways are specifically undergoing plastic changes after chronic SCI and contribute to the loss of presynaptic inhibition associated with hyperreflexia. While we can confirm with certainty that presynaptic inhibition initiated by PBSt on the plantar H-reflex is disrupted by chronic SCI, whether the decreased transmission in PBSt PAD pathway is the consequence of a change in interneuronal excitability,failure to activate said interneurons by sensory afferents and/or a decrease in PAD activation remains to be determined.

### Step-training restores transmission in PBSt group I PAD pathways after chronic SCI

Our results indicate that step-training did not affect input-output curve of L4-DRPs evoked by the tibial nerve suggesting that this pathway is not undergoing activity-dependent plasticity after SCI. In contrast, we found that step-training restored the amplitude of L4-DRPs evoked by PBSt group I muscle afferents, which prevented the shift toward a requirement for higher intensity of stimulation to evoke a DRP_max_. Tib-DRPs were barely affected by chronic SCI, while PBSt-DRPs were significantly decreased, suggesting that step-training may specifically target pathways that are disrupted by chronic SCI. Together, these results suggest that the SCI-induced decrease in transmission in PAD pathways activated by PBSt group I afferents is altered in an activity-dependent manner, whereas PAD pathways activated by PBSt group II or tibial cutaneous afferents (historically termed flexor reflex afferents) are not. However, cutaneous afferents can evoke different PAD patterns in similarly identified afferents, including a decrease in PAD evoked by PBSt group I afferents (Quevedo *et al.*, 1993; 1995), the occurrence of which can be shifted after a neuronal insult such as a crush injury (Enriquez *et al.*, 1996a). Whether chronic SCI or step-training altered the PAD pattern evoked by tibial cutaneous afferents remains to be determined as it would not have been reflected by Tib-DRPs input-output properties.

### Presynaptic and postsynaptic inhibition of the H-reflex are regulated in an activity-dependent manner

Our data show that step-training did not modify the time course, but noticeably increased the strength of presynaptic inhibition. After chronic SCI and other conditions inducing long-lasting immobilization, motor training decreases the hyperexcitability of the monosynaptic reflex and improves reflex modulation (Côté *et al.*, 2003; Côté & Gossard, 2004; Martin Ginis & Latimer, 2007; Lundbye-Jensen & Nielsen, 2008b; Knikou & Mummidisetty, 2014; Caron *et al.*, 2016). Accordingly, our results show that the slope of the plantar H-reflex recruitment curve is increased after chronic SCI, suggesting a greater recruitment of motor pools by tibial Ia afferents (Capaday, 1997). This shift was not observed in step-trained animals, in agreement with previous studies suggesting that step-training decreases the slope of the H-reflex recruitment curve in SCI patients (Smith *et al.*, 2015) and more generally that the slope of the H-reflex recruitment curve is altered in an activity-dependent manner. (Lundbye-Jensen & Nielsen, 2008a). A decreased recruitment of motor pools suggests a less homogenous activation of motor units.

SCI and immobilization induce changes in the intrinsic properties of slow motor units towards faster motor unit properties (Burnham *et al.*, 1997; Talmadge, 2000; Beaumont *et al.*, 2004; Cormery *et al.*, 2005). Motor activity can counteract these effects by shifting back fast motor unit properties towards slower properties (Yarar-Fisher *et al.*, 2018). Although not directly measured in this study, our results suggest a possible contribution of an activity-dependent alteration in motor unit properties to the normalization of the motor pool recruitment and to the excitability of the H-reflex. In addition to a return in the recruitment gain of the plantar H-reflex with step-training, our results show that both short-(10-20ms C-T interval) and long-lasting inhibition (50-150ms C-T intervals) induced by PBSt group I afferents are restored with step-training, suggesting a return of presynaptic inhibition of the plantar H-reflex, and a likely restoration of postsynaptic inhibition. A return of presynaptic inhibition of the H-reflex with locomotor training has been suggested in extensor muscles both in animals (Côté *et al.*, 2003) and humans (Knikou & Mummidisetty, 2014). The increase in DRP amplitude evoked by PBSt group I afferents in step-trained animals suggests that the recovery of presynaptic inhibition of the plantar H-reflex relies on a larger PAD evoked by PBSt on tibial Ia afferents. While our results suggest a return of postsynaptic inhibition, most likely through increased IPSPs in motoneurons also evoked by PBSt group I afferents (Hultborn *et al.*, 1987a; Stuart & Redman, 1992), it remains to be demonstrated. Together, our results suggest that the repetitive activation of sensory afferents triggered by step-training decreases hyperreflexia through a simultaneous normalization in the recruitment of motor pools and a restoration in both presynaptic and postsynaptic inhibition induced by PBSt group I afferents.

### Possible mechanisms contributing to the reorganization of afferent control of spinal networks after SCI and step-training

In this study, we stimulated PBSt, which is an efficient sensory source of presynaptic inhibition frequently used to inhibit the monosynaptic reflex (Frank & Fuortes, 1957; Eccles *et al.*, 1962c; Côté & Gossard, 2003). Presynaptic inhibition is largely mediated by the shunting action of GABA_A_ receptors at the afferent terminal, which decreases the size of the action potential that invades the terminal and reduces the activation of calcium current and associated transmitter release (Curtis *et al.*, 1971; Stuart & Redman, 1992; Curtis, 1998; Cattaert & El Manira, 1999). This inhibition is described as the *phasic* PAD, in opposition to the *tonic* PAD, which regulates afferents transmission through extrasynaptic GABA_A_ receptors (Lucas-Osma *et al.*, 2018). While a change in tonic PAD has the potential to be of particular importance for the propagation of action potentials at branching points, our recording configuration for DRPs (6mm interelectrode distance and 0.1Hz high pass) prevented its estimation in this experiment. Overall, whether the decreased transmission in PBSt PAD pathway is the consequence of a change in interneuronal excitability, failure to activate said interneurons by sensory afferents and/or a decrease in *phasic vs. tonic* PAD activation on primary afferents remains to be determined.

More recently, last order GABAergic interneurons specifically involved in presynaptic inhibition have been genetically identified in rodent models (Fink *et al.*, 2014; Koch *et al.*, 2017) and presynaptic inhibitory mechanisms were shown to rely not only on GABAergic signaling, but also on other neurotransmitters, neuromodulators and receptors that also could contribute to our results (Kremer & Lev-Tov, 1998; Hochman *et al.*, 2010; Lucas-Osma *et al.*, 2018). Finally, mechanisms other than PAD are also likely to contribute to presynaptic inhibition of Ia afferents and possibly contribute to the PAD-independent presynaptic inhibition of the H-reflex in our experiments (Price *et al.*, 1984; Stuart & Redman, 1992; Hultborn *et al.*, 1996; Curtis *et al.*, 1997; Garcia-Ramirez *et al.*, 2014).

#### Clinical significance

Pathological changes in presynaptic inhibition are crucial to several diseases including peripheral inflammation, abnormal pain processing, hyperreflexia and spasticity (Wall & Devor, 1981; Calancie *et al.*, 1993; Enriquez-Denton *et al.*, 2004; Witschi *et al.*, 2011). Sensory feedback plays a critical role in determining and refining spatiotemporal features of motor output (Rossignol *et al.*, 2006). The spinal cord is continuously flooded by sensory input via primary afferents, and the integrity of presynaptic inhibitory mechanisms to effectively select the relevant information and filter sensory noise is critical.

The CNS is believed to adjust the level of presynaptic inhibition of extensor group Ia afferents to modulate the central gain of the stretch reflex (Capaday & Stein, 1986; Hultborn *et al.*, 1987b), and also the locomotor rhythm in animals and humans (Simonsen & Dyhre-Poulsen, 1999; Morita *et al.*, 2001; Knikou, 2006). For example, it was shown that maintaining the Achilles’ tendon stretched maintains extensor muscles activity, which during locomotion prevents the switch from stance to swing phase (Duysens & Pearson, 1980). Thus, presynaptic inhibition evoked by PBSt group I afferents could play an important role in the initiation of the swing phase when the Achilles’ tendon is loaded during stance. In contrast, plantar cutaneous afferents inhibit presynaptic inhibition evoked by PBSt group I afferents on Ia afferents (Quevedo *et al.*, 1993; 1995), which during locomotion could contribute to increase extensor muscles activity and decrease the inhibition of the extensor H-reflex with ground contact (Duysens & Pearson, 1976; Duysens, 1977; Guertin *et al.*, 1995; Iles, 1996). Thus, the decrease in presynaptic inhibition observed after chronic SCI impairs this ability to tailor motor output to environmental constraints and may contribute to the presence of co-contraction during walking preventing the transition from stance to swing and to the development of tremors or clonus, which are frequently observed in spastic patients (Nielsen *et al.*, 2007).

The recovery of presynaptic inhibition with step-training illustrates the importance of activity-based therapies after chronic SCI. Understanding the physiological effects of rehabilitation programs is critical to the design of optimized therapies in the clinic. For example, step-training changes the pattern of postsynaptic potentials evoked by cutaneous afferents in specific motor pools. It more specifically leads to an increase in the excitation of ankle extensors by cutaneous afferents from the plantar surface of the foot, presumably to improve weight-bearing during ground contact (Côté & Gossard, 2004; Knikou, 2010). The concomitant increase in presynaptic inhibition evoked by PBSt group I afferents would not only counteract the aforementioned debilitative effects of decreased presynaptic inhibition after SCI, but would also allow for the modulation of sensory inputs resulting from plastic changes after SCI. As presynaptic inhibition is regulated in a task-dependent manner (Côté & Gossard, 2003), more studies are needed to investigate if the restoration of presynaptic inhibition is specific to certain movements and how it responds to a comprehensive rehabilitation program that includes an array of motor training and pharmacological tools to manage spastic symptoms.

## Additional information

### Authors contributions

All experiments were performed in the Côté Lab at the Marion Murray Spinal Cord Research Center at Drexel University College of Medicine, Philadelphia, Pennsylvania, USA. GC and MPC were responsible for the conception and design of the work. GC and JB performed the experiments. GC performed the analysis. GC and MPC interpreted the results. GC drafted the manuscripts. All authors edited and revised the manuscript and approved the final version for publication. MPC is responsible for all aspect of the reported work, including the integrity of the data collected and the accuracy of data analysis.

#### Funding

This work was supported by grants from the National Institute of Neurological Disorders and Stroke (RO1 NS083666) and the Craig H. Neilsen Foundation (189758).

## Acknowledgements

We thank Dr. Jean-Pierre Gossard for helpful comments on an earlier version of this manuscript and Kyle Yeakle for assistance in data collection.

